# Selection and the direction of phenotypic evolution

**DOI:** 10.1101/2022.05.28.493855

**Authors:** François Mallard, Bruno Afonso, Henrique Teotónio

## Abstract

Predicting phenotypic evolution on the short-term of tens to hundreds of generations, particularly in changing environments and under finite population sizes, is an important theoretical goal. Because organisms are not simply collections of independent traits, making headway into this goal requires understanding if the phenotypic plasticity of ancestral populations aligns with the phenotypic dimensions that contain more genetic variation for selection to be effective and eventually feedback on the maintenance of genetic variation and promote adaptation or rescue from extinction. By performing 50 generations of experimental evolution in a changing environment we show that ancestral phenotypic plasticity for locomotion behavior in the partially-outcrossing nematode *Caenorhabditis elegans* is nonadaptive because it does not align with the phenotypic dimension encompassing most genetic variance in the ancestral population and is of no consequence to future phenotypic divergence. Despite evolution of the genetic structure of locomotion behavior we are able to predict the direction of phenotypic divergence, but not the magnitude, based on the genetic covariances between the component traits of locomotion behavior and fitness of the ancestral population. We further demonstrate that indirect selection on the component traits of locomotion behavior with unobserved trait(s) is responsible for the observed phenotypic divergence on them. Our findings indicate that selection theory can predict the direction of short-term adaptive phenotypic evolution.

## 2 Introduction

Phenotypic evolution should be considered in the context of the whole organism, as organisms are not just collections of independent traits (Gould and Lewontin, 1979). Theory indicates that stabilizing and disruptive selection on each trait, and correlated selection between trait combinations, which together define the individual “selection surface” (Phillips and Arnold, 1989), will shape standing additive genetic variances of traits and covariances between traits (Arnold et al., 2001; Jones et al., 2004; Lande, 1980; Svensson et al., 2021), as summarized by the **G** matrix describing the traits’ genetic structure that is inherited between parents and offspring. This is because the phenotypic dimensions with greater additive genetic variation allow less constrained responses to directional selection and thus more rapid adaptation to novel environments or rescue from extinction (Blows and McGuigan, 2015; Chevin et al., 2010; Lande, 2009; Schluter, 1996). Theory also suggests that *de novo* input of genetic covariances by pleiotropic mutations, if variable, could in the long-term of mutation-selection balance (time being scaled by the effective population size), be aligned with the orientations of the selection surface and eventually explain phenotypic divergence between populations and species (Jones et al., 2007, 2014). However, many studies of natural populations find more standing genetic variation than that expected at mutation-selection balance (Sella and Barton, 2019; Walsh and Lynch, 2018), and it is not entirely clear that the **G** matrix can evolve to align with the selection surface (Arnold et al., 2008; Steppan et al., 2002), in part because selection might not be constant or uniform when populations are faced with changing environments (Careau et al., 2015; Chevin and Haller, 2014; Gomulkiewicz and Houle, 2009; Guzella et al., 2018), in part because all populations are finite, mutations rare, founder effects common, and drift on the **G** matrix likely to alter phenotypic evolution on the short-term of tens to hundreds of generations, before mutation-selection balance is reached (Mallard et al., 2022, 2019; Matuszewski et al., 2015; Phillips et al., 2001; Whitlock et al., 2002).

When populations are challenged in a novel environment it is possible that phenotypic plasticity, the ability of a genotypes to express alternative trait values under environmental variation, aids immediate population persistence (Ghalambor et al., 2007; Pfennig et al., 2010), but for adaptive phenotypic divergence to then occur the plastic phenotypic dimension should align with the dimension where most genetic variation is found (Draghi and Whitlock, 2012; Gomulkiewicz and Holt, 1995; Price et al., 2003; Via and Lande, 1985). Adaptation or rescue from extinction can be hampered if less genetic variation is available for selection in the relevant phenotypic dimensions. A recent meta-analysis of natural populations has found that phenotypic dimensions of plasticity and genetic variation are aligned (Noble et al., 2019), despite great heterogeneity among the studies surveyed, suggesting that the development and physiology responsible for the traits’ expression under environmental variation reflect the distribution of pleiotropic mutational effects due to past correlated selection (Draghi and Whitlock, 2012; Jones et al., 2014; Morrissey, 2015). Yet, several studies found that the phenotypic plasticity of ancestral populations does not always match the phenotypic directions of adaptation (Ghalambor et al., 2015; Ho and Zhang, 2019; Koch and Guillaume, 2020; Mallard et al., 2020; Sikkink et al., 2019), indicating that selection is often misunderstood.

While accounting for phenotypic plasticity is relatively straightforward in lab populations, or natural populations that can be transplanted between environments, the same cannot be said about understanding selection. Estimates of selection will be biased in strength or kind if unobserved traits impacting fitness are genetically correlated with the observed traits or there is correlated selection because of environmental covariances between observed and unobserved traits with fitness, both instances of “indirect” selection on the observed traits (Morrissey et al., 2010; Rausher, 1992; Walsh and Lynch, 2018). Lande’s equation (Lande, 1979), 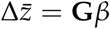, models trait evolution over one generation in the multivariate context 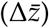 as a function of the **G** matrix and the directional selection gradients on each trait (*β*), while explicitly considering potential correlated selection between the observed traits and stabilizing/disruptive selection on each one of them (Lande and Arnold, 1983), but might fail when unobserved traits are important. Using the Robertson secondary theorem of natural selection (Robertson, 1968; Walsh and Lynch, 2018), 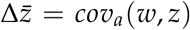, however, observed mean trait changes over one generation are accurately predicted because the responses due to selection, i.e., the selection differentials, equal the breeding value change of the trait affecting fitness and thus the additive genetic covariance of the trait with relative fitness (w), independently of unobserved traits (Morrissey and Bonnet, 2019; Morrissey et al., 2010). However, distinguishing direct from indirect selection on each trait is not possible with the Robertson equation, and further the parental trait breeding values must have no transmission bias, due for example to non-Mendelian segregation or to transgenerational environmental effects, and be fully mapped onto offspring trait breeding values (Price, 1972; Walsh and Lynch, 2018).

To help resolve these problems, and following Etterson and Shaw (2001) and Morrissey et al. (2012) among others, Stinchcombe et al. (2014) have proposed in a single statistical framework to apply the Robertson equation in a presumed ancestral population, before phenotypic divergence, to predict the direction of trait evolution because of selection, and in a second step to apply Lande’s equation to infer the gradients of directional selection on each trait: 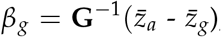; where the selection differential *s*, or the trait response *R* because of selection after one generation, is defined by the difference between the trait means for an observed ancestral population 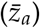 and the expected trait means of a divergent population as predicted by the Robertson equation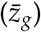. For example, if the “genetic” selection gradient during the period considered (*β*_*g*_) is not inferred, or is different from the Lande’s “phenotypic” selection gradient obtained by the regression of divergent onto ancestral trait values (*β*), then phenotypic divergence must have been due to indirect selection. This framework is appropriate for a single generation and it is unknown if it can be applied to multiple generations and changing selection surfaces. Furthermore, as the **G** matrix evolves by selection, drift or mutation, the prediction of phenotypic divergence from the Robertson equation and the inference of selection with Lande’s equation likely becomes inaccurate (Phillips and McGuigan, 2006; Shaw et al., 1995), though we lack expectations of when they become so, except for models at mutation-selection balance that notably disregard drift, strong selection or reduced effective recombination between fitness loci (Arnold et al., 2001; Lande, 1980; Mallard et al., 2022, 2019; Sella and Barton, 2019).

We here report a test of Stinchcombe et al. (2014) proposal by doing experimental evolution from standing genetic variation in the partially-outcrossing nematode *Caenorhabditis elegans* and following as a phenotype the worms’ locomotion behaviour. Locomotion behavior is described by six statistically independent traits of transition rates between movement state and direction (Mallard et al., 2019), measured in a common scale with no need for transformation to apply selection theory (Houle et al., 2011). We started by characterizing the broad-sense **G** matrices of a lab population in the low salt concentration environment where it had adapted to for 140 generations (Chelo and Teotónio, 2013; Noble et al., 2017; Teotónio et al., 2012), as well as in a novel high salt concentration environment, by measuring locomotion differences between inbred lines derived from the outbreeding population. We have previously shown that the broad-sense **G** matrix is an adequate surrogate of the true narrow-sense additive **G** matrix of experimental outbreeding populations (Mallard et al., 2019), but see (Chelo et al., 2019) and Discussion. We then challenged the lab adapted population to a gradually increasing salt concentration environment for 35 generations, followed by 15 generations in constant high salt and measured locomotion behavior in three derived replicate populations. We also before showed that derived populations adapted to the high salt conditions (Theologidis et al., 2014), and thus here ask if the Robertson equation, when applied to the ancestral lab adapted population, predicts 50 generations of locomotion behavior evolution in the novel high and ancestral low salt environments, and whether direct and indirect selection have a bearing on the observed transition rate divergence in the target high salt environment, using Lande’s equation.

## 3 Methods

### 3.1 Archiving

Fertility data (see below) has been previously published in Chelo et al. (2019). New data, R code for analysis and modeling results (e.g. **G** matrix estimates) can be found in our github repository and will be archived in *Dryad*.*org* upon publication.

### 3.2 Experimental populations

Three replicate populations were derived from an outbreeding population (named A6140; “A” for androdioecious, “6” for replicate six, “140” for generations of lab evolution), previously adapted to standard lab conditions (Noble et al., 2017; Teotónio et al., 2012; Theologidis et al., 2014). A6140 in turn resulted from the hybridization of 16 founder strains during 33 generations followed by 140 4-day discrete and non-overlapping life-cycles at *N* = 10^4^ census sizes at the time of reproduction (Teotónio et al., 2012), and *N*_*e*_ = 10^3^ effective population sizes (Chelo and Teotónio, 2013). *C. elegans* is an androdioecious roundworm, where hermaphrodites can self but only outcross when mated with males. Natural populations are depauperate of genetic diversity and males are rare due to a long history of selfing, selective sweeps and background selection (Andersen et al., 2012; Rockman et al., 2010). Under our lab conditions, however, partial-outcrossing is readily maintained at frequencies between 60%-80% (Mallard et al., 2019; Teotónio et al., 2012). During domestication populations were maintained in 10x 9cm Petri dishes NGM-lite agar media containing 25mM NaCl and a lawn of *E. coli* HT115 that served as food from the L1 larval stage until reproduction. Periodic population samples were cryogenetically stored (Stiernagle, 1999), allowing for contemporaneous comparisons between ancestral and derived populations after revival of samples (Teotónio et al., 2017).

Here we report the evolution of locomotion behavior in three replicate populations (named GA[1,2,4] populations: “G” for gradual, “A” for androdioecious, “1,2,4” for replicate number) derived from A6140. GA populations were maintained in exactly the same conditions as during domestication except that the NGM-lite media was supplemented with 8mM of NaCl at each generation for 35 generations and then kept constant at 305mM NaCl for an additional 15 generations. Details about the derivation of the GA populations can be found in Theologidis et al. (2014). In Theologidis et al. (2014) we showed, using competitions assays between ancestral and derived populations, that there was adaptation and further in Chelo et al. (2019) that there was maintenance of partial-outcrossing and heterozygosity of about 0.25 for 1 single nucleotide variant per kbp by the end of the 50 generations of evolution. Control populations were also kept in the domestication environment and the evolution of locomotion behavior in these populations is described in a (Mallard et al., 2019). Throughout we refer to the NGM-lite 305mM NaCl environment as the “high” salt target environment, while the domestication 25mM NaCl environment as the “low” salt environment.

### 3.3 Inbred lines and fitness

We derived inbred lines by selfing single hermaphrodites from the ancestor (A6140) and the three replicate populations at generation 50 (GA[1,2,4]50) for at minimum of 10 generations (Chelo et al., 2019). The total number of lines obtained and phenotyped for locomotion behavior can be found in Table 1. In Chelo et al. (2019) we measured individual hermaphrodite self-fertility in high salt, a trait that here we use as a surrogate for fitness. Fertility was measured under environmental conditions that closely followed those of experimental evolution, except for density and presence of males. The log-transformed, covariate-adjusted fertility values (best linear unbiased estimates) for each inbred line were downloaded from Chelo et al. (2019), exponentiated, and divided by the mean to obtain a relative fitness measure (*w*).

**Table 1:** Number of measurements per population in each of three assay years. In total there were 188 lines for A6140 and 61, 61 and 42 lines for GA[1,2,4].

| Assay Year | Low Salt |  |  |  | High Salt |  |  |  |
| --- | --- | --- | --- | --- | --- | --- | --- | --- |
|  | Ancestral | Generation 50 |  |  | Ancestral | Generation 50 |  |  |
|  | A6140 | GA150 | GA250 | GA450 | A6140 | GA150 | GA250 | GA450 |
| 2012 | 167 | 70 | 58 | 53 | 169 | 73 | 58 | 53 |
| 2013 | 194 | 44 | 58 | 29 | 191 | 44 | 59 | 30 |
| 2014 | 0 | 0 | 0 | 0 | 8 | 2 | 4 | 0 |

### 3.4 Locomotion behavior

Inbred lines were thawed from frozen stocks on 9 cm Petri plates and grown until exhaustion of food. This occurred 2-3 generations after thawing, after which individuals were washed adults removed by centrifugation, and three plates per line seeded with 1000 larvae at mixed larval stages. Samples were then maintained for one to two complete generations in the standard domestication environment. At the assay generation (generation 4-6 generations post-thaw), starvation-synchronized L1 larvae were seeded in both the low and high-salt environments. Adults were phenotyped for locomotion behaviour 48h later at their usual time of reproduction in one 9cm plate. At the beginning of each assay we measured ambient temperature and humidity in the imaging room. Given the number of lines phenotyped we repeated this protocol several times by the same experimenter, each defining a statistical “block”, usually including samples from A6140 and from one of the GA populations, over a period of three years (Table 1). In total, we phenotyped 188 lines from the A6140 population and 61,61 and 42 lines from each of the GA[1,2,4] populations, with most lines being phenotyped twice and always in separate blocks (average 1.9 in low salt, 2 in high salt).

We imaged adult hermaphrodites using the Multi-Worm Tracker [version 1.3.0; Swierczek et al. (2011)], and followed the protocols detailed in Mallard et al. (2019) to collect the data. Each movie contains about 1000 tracks of hermaphrodites (called objects) with a mean duration of about 1 minute. Standardized to a common frame rate, we filtered and extracted the number and persistence of tracked objects per movie, and assigned movement states across consecutive frames as forward, still or backwards (assuming forward as the dominant direction of movement).

Modeling transitions rates has also been detailed in Mallard et al. (2019). Briefly, expected transition rates between forward, still and backward movement states follow a continuous time Markov process, and given the constraint that self-transition rates equal the sum of the six non-self transition rates, we considered that these non-self transition rates are independent [called *q*_*k*_ for notation consistency with Mallard et al. (2019)]. For estimation we used log-likehood models specified in RStan [Stan Development Team (2018), R version 3.3.2, RStan version 2.15.1], which performs Bayesian inference to calculate the posterior probability of the parameters’ distribution given the observed data.

### 3.5 Phenotypic plasticity and divergence

We modeled phenotypic plasticity among high-and low-salt environments in A6140 and GA[1,2,4]50 using linear mixed models. For each transition rate:

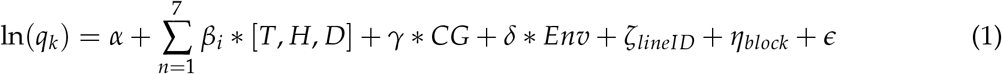

where *α* is the overall mean, *β*_*i*_ the set of environmental fixed effects accounting for temperature, humidity and density (including interactions), *γ* is a fixed effect accounting for mean differences between assay years, 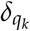 the mean difference between our two environments, *ζ* ∼ 𝒩 (0, *σ*^2^) and *η* ∼ 𝒩 (0, *σ*^2^) the random effects of line identities and block effects, respectively, and finally, ∼ 𝒩 (0, *σ*^2^) the residual error. Each measured variable was centered and standardized prior to the analysis.

We modeled divergence of locomotion behavior after 50 generations of evolution using two different formulations. In the first (equation 2), we tested for a consistent divergence among replicates by considering GA[1,2,4]50 identity as a random effect. In the second (equation 3), we further investigated individual replicate divergence from A6140.

For the first model:

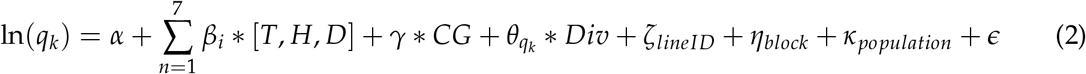

with *α, β*_*i*_, *γ, δ, ζ, η* and as defined above. *θ*_*k*_ is here the mean difference between the ancestral and the evolved populations. *κ* ∼ N (0, *σ*^2^) is a random effect of mean differences between our three GA[1,2,4] evolved populations.

For the second model:

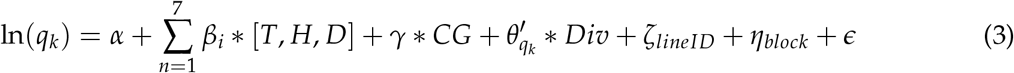

where 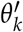 is now a fixed categorical effect accounting for mean differences between our populations. Note that the *κ* random effect does not appear in this model.

The significance of fixed effects were tested with Likelihood Ratio Tests (LRT, approximately *χ*^2^ distributed), using the *anova* command in R with arguments being the two nested models containing or not the fixed effect being tested. Differences between A6140 and each of the GA replicates were tested using the *emmeans* package (Lenth, 2021).

We define from equations 1 and 2 the vectors of phenotypic plasticity Δ*P* and of phenotypic divergence Δ*D* as:

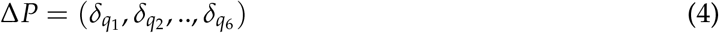

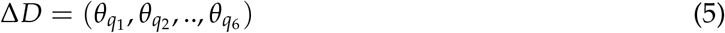

Note that there is one Δ*D* for each environment. We further created distributions of Δ*D* and

Δ*P* by bootstrapping 1000 times the set of inbred lines before estimating each of the vectors. These are used to calculate the confidence intervals of Figures 3, 5, 6, S6, and S8.

### 3.6 G matrices

We estimated **G** matrices separately for each of the four studied populations (A6140, GA[1,2,4]50). We present results for the A6140 population in the low and high salt environment, for GA[1,2,4]50 only in the high salt environment. The 6 transition rates *q*_*k*_ were fitted as a multivariate response variable *y* in the model:

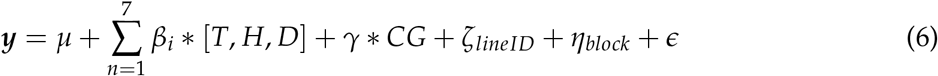

here *µ* is the intercept and *β*_*i*_, *γ, ζ, η* and as defined above (equation 1). The matrix of genetic (co)variances is then half the line (co)variance matrix (*ζ*_*lineID*_), as we have measured inbred lines we twice count genetic effects. We have elsewhere showed that selection is significant during inbreeding (Chelo et al., 2019), but we have also shown that it does not on average change the mean transition rate values in low salt (Mallard et al., 2019). The broad-sense **G** matrix as estimated here should thus be an adequate surrogate for the narrow-sense **G** matrix (Lynch and Walsh, 1998), and see Discussion.

Models were fit with the R package *MCMCglmm* (Hadfield, 2010). We constructed priors as the matrix of phenotypic variances for each trait. Model convergence was verified by visual inspection of the posterior distributions and an autocorrelation below 0.05. 100,000 burn-in iterations were done with a thinning interval of 2,000 and a total of over 2 million MCMC iterations.

We define the main axis of genetic variation *g*_*max*_ as the first eigenvector of our A6140 **G**-matrix, *g*_2−6_ its subsequent eigenvectors and *λ*_1−6_ their six eigenvalues. To describe the genetic dimensions of divergence among all the four populations (A6140 and GA[1,2,4]50), we performed spectral decomposition of the four **G** matrices in high salt; for details see Aguirre et al. (2014); Morrissey and Bonnet (2019). **G**-matrix differences in this analysis are described by orthogonal eigentensors, which in turn can be decomposed into eigenvectors with associated eigenvalues. We define the first of these eigenvectors as the genetic divergence vector *e*_11_ as it is the phenotypic dimension encompassing most of the genetic differences between all four **G** matrices.

As discussed in Walter et al. (2018), the significance of the posterior mean variance-covariance estimate is based on the overlap between the posterior null distribution of the posterior mean with the observed posterior mean. We consider that differences between the estimated empirical distributions can be inferred when their 80% credible intervals (strickly 83%) do not overlap (Austin and Hux, 2002).

### 3.7 Selection

We also estimated *G*_*qw*_ matrices, as defined by Stinchcombe et al. (2014), with 13 traits: the six transition rates estimated in each environment (*q*_*k*_) and the relative fertility measured in the high salt environment (*w*). By computing *G*_*qw*_ we estimate the (broad-sense) genetic covariance between transition rates and relative fertility. According to Robertson’s secondary theorem of natural selection (Robertson, 1968; Walsh and Lynch, 2018) if the broad-sense genetic covariance is the same as the narrow-sense additive genetic covariance between traits and fitness, then it will be equal to the selection differentials on each transition rate, *s*, and thus the responses to selection *R*:

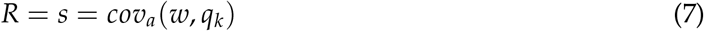

Following Stinchcombe et al. (2014), we then estimated the “genetic” directional selection gradients on each transition rate *β*_*g*_ using Lande’s equation:

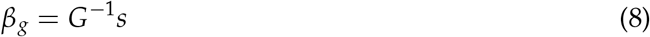

Computation was done as described above for the **G** matrices, with the same parameters.

### 3.8 Plasticity, genetic variances, selection and divergence

Our analysis relies on comparing the direction or the size/length of several vectors and scalars in multivariate space. We used the metrics introduced by Noble et al. (2019) to compare the alignment of the phenotypic dimension with most genetic variation, *g*_*max*_, with phenotypic plasticity, Δ*P*, in A6140. The angle between *g*_*max*_ and Δ*P* is:

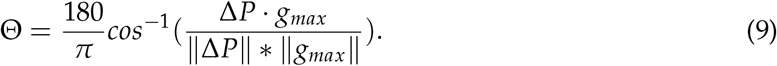

As both *g*_*max*_ and −*g*_*max*_ are the first eigenvector of the **G-**matrix and their eigenvalue 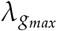 a single (positive) scalar, Θ values between 90° and 180° were transformed so that Θ always remains between 0° and 90° (Θ = 180°-Θ, which results from using −*g*_*max*_ instead of *g*_*max*_ in equation 9). The null expectation for Θ when using random vectors is then 45°.

To estimate the amount of genetic variance in plasticity:

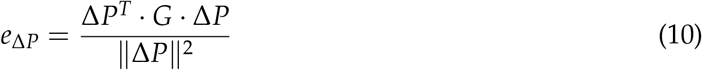

Π being next defined as the ratio between the genetic variance along plasticity dimensions and the maximum amount of variance in any phenotypic direction (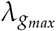, the first eigenvalue of the **G** matrix):

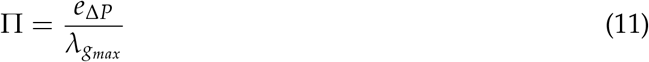

Π values are comprised between 0 (no genetic variance along the plasticity dimensions) and 1 (when plasticity contains all genetic variance). The null expectation of Π being:

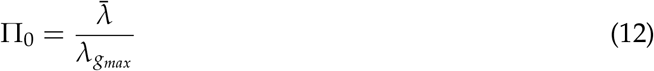

where 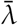 is the mean of all six eigenvalues of **G**, in turn the average genetic variance across any phenotypic dimensions (i.e.,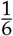 of the total genetic variance).

We defined other angles to compare the alignment of ancestral *g*_*max*_ with the dimensions of phenotypic and genetic divergence, by replacing in equation 9 Δ*P* with Δ*D* and *e*_11_, respectively. We also computed the angle between ancestral plasticity Δ*P* and phenotypic divergence Δ*D*, and that between predicted selection responses (*R*/*s* from equation 7) with observed divergence (Δ*D*) as a test of Robertson’s equation. For all metrics, we computed 80% and 95% credible intervals by randomly sampling the posterior distribution of the A6140 **G** matrix, the exception being the angle between plasticity and phenotypic divergence, where we used the least square estimates from equations 1 and 2.

## 4 Results

### 4.1 Locomotion behavior in the ancestral population

Our ancestral population (A6140) evolved for 140 generations in a defined lab environment characterized in part by low salt (25mM NaCl) in the worms’ growing media (see Methods). We found significant phenotypic plasticity for locomotion behavior between the low salt environment and a novel high salt (305mM NaCl) environment (Figure 1). Only for one of the six transition rates measured to describe locomotion behavior, the transition rate between backward and forward movement did not show plasticity (Table 2). Three transition rates increased (still to/from forward and backward to still) and two decreased (still/forward to backward) in high salt when compared with low salt.

**Figure 1:**
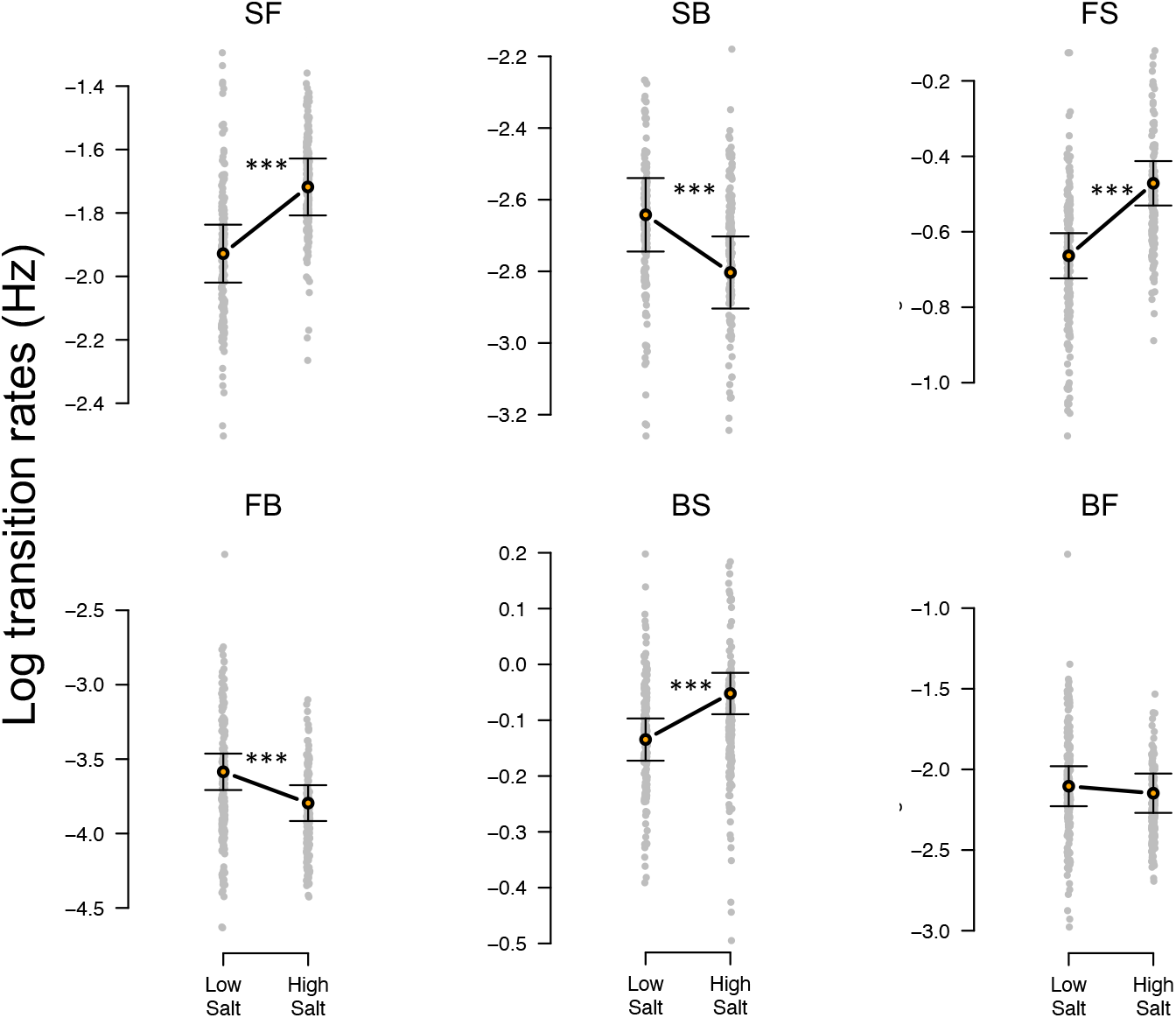
Phenotypic plasticity in the ancestral population A6140. Grey dots indicate the values estimated for each inbred line in the low and high salt environments for each of the transition rates between activity and movement direction. F stands for “forward”, B for “backward” and S for “still”, left to right order indicating movement direction. Orange shows the mean least-square estimates of the transition rates among inbred lines in each environment, with bars the 95% confidence intervals (see Methods). Mean differences between environments are indicated by a line, with asterisks showing plasticity (LRT *χ*^2^, P-value <0.001; Table 1).

**Table 2:** Phenotypic plasticity in the ancestral population

| transition rate | mean low salt | plasticity | Chisq | df | P value |
| --- | --- | --- | --- | --- | --- |
| SF | -1.7 | 0.21 | 25.2 | 1 | 5.1 e-07 |
| SB | -2.8 | -0.16 | 11.5 | 1 | 6.9 e-04 |
| FS | -0.47 | 0.19 | 43.4 | 1 | 4.4 e-11 |
| FB | -3.8 | -0.21 | 13.1 | 1 | 2.8 e-04 |
| BS | -0.052 | 0.082 | 18.8 | 1 | 1.5 e-05 |
| BF | -2.1 | -0.043 | 0.579 | 1 | 0.45 |

We next partitioned the phenotypic (co)variances of our six transition rates among the inbred lines of A6140 into genetic and error (co)variances in each salt environment. We find that the structure of the two **G** matrices is quite similar (Figure 2). First, there is no detectable difference in the total genetic variance of the **G** matrices (Figure 2A). Second, in both environments, locomotion behavior is modular (Figure 2B). Transition rates from still to forward or backward, i.e. leaving the still state, are negatively correlated with the remaining transition rates and these positively correlated with each other. Third, eigen decomposition of the **G** matrix in low salt shows that it is rounder than that of high salt, but differences are not important (Figure 2C). The first eigenvector dimension (*g*_*max*_) encompasses most genetic variation in both environments, and they are aligned as the angle between them is small (mean ± 95 CI: 12.6^°^[4.4 − 21.6]).

**Figure 2:**
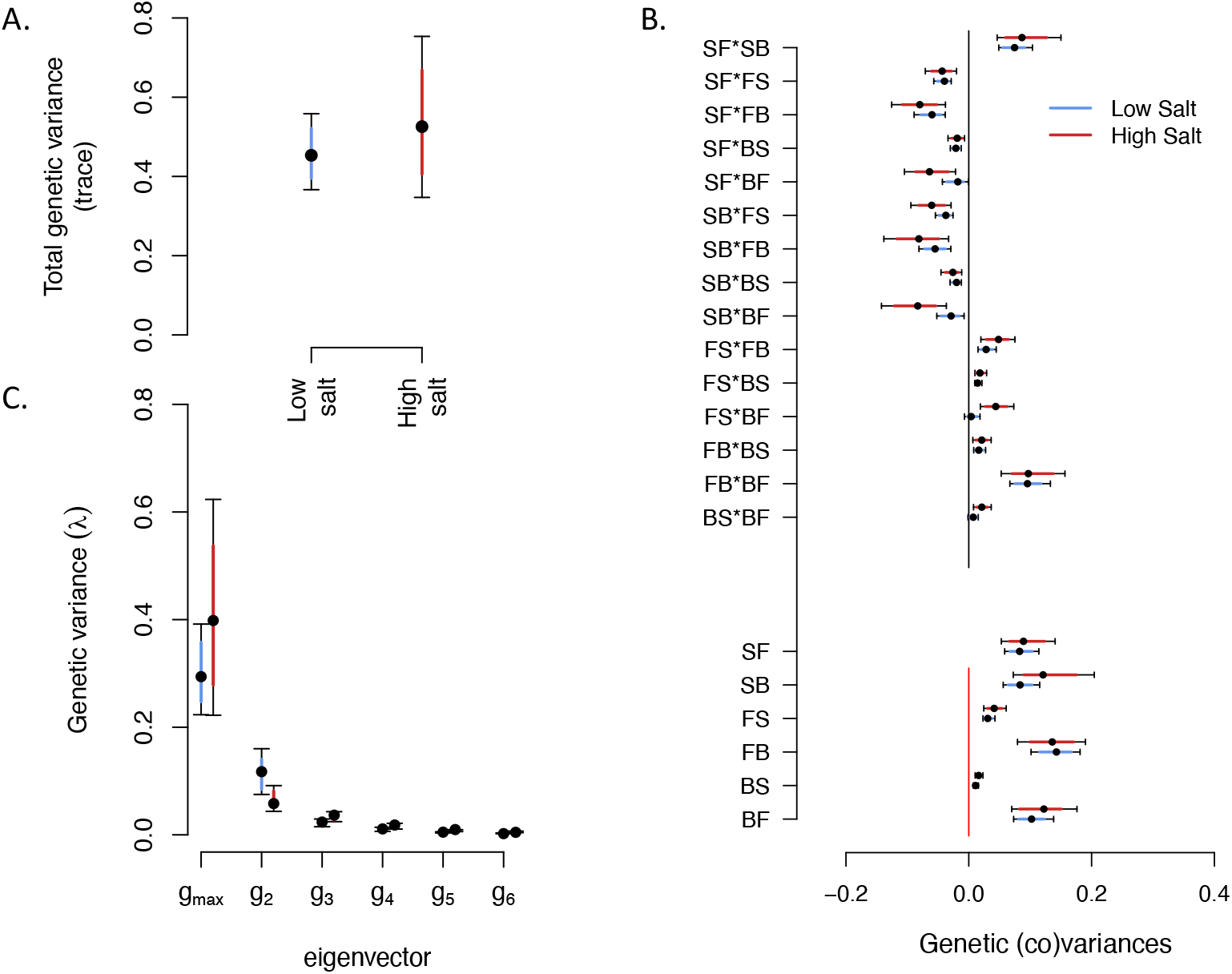
**G**-matrix of the ancestral population A6140 measured in low and high salt environments. **A**. Total genetic variance in each environment (trace of the **G** matrices), blue (red) for low (high) salt environments. **B**. Bottom six entries indicate the diagonal estimates the genetic variances of transition rates, top 15 entries indicate the off-diagonal estimates of genetic covariances between transition rates. Blue (red) indicate estimates in low (high) salt, with colored intervals and bars showing respectively 80% and 95% credible intervals of the posterior distribution, dots the mean of the posterior (see Methods). In Supplementary Figure S1, we show that most (co)variance estimates, in both environments, are different from those expected just with sampling effects. **C**. Eigenvalues estimates of the six eigenvectors of the **G** matrices, blue (red) for low (high) salt. Dot and bars indicate the mean, 80% and 95% credible intervals of the posterior distributions. Following Morrissey and Bonnet (2019), the two phenotypic dimensions with most genetic variation, *g*_*max*_ and *g*_2_, in low salt, but only *g*_*max*_ in high salt, are different from null expectations in both environments (not shown).

**Figure 3:**
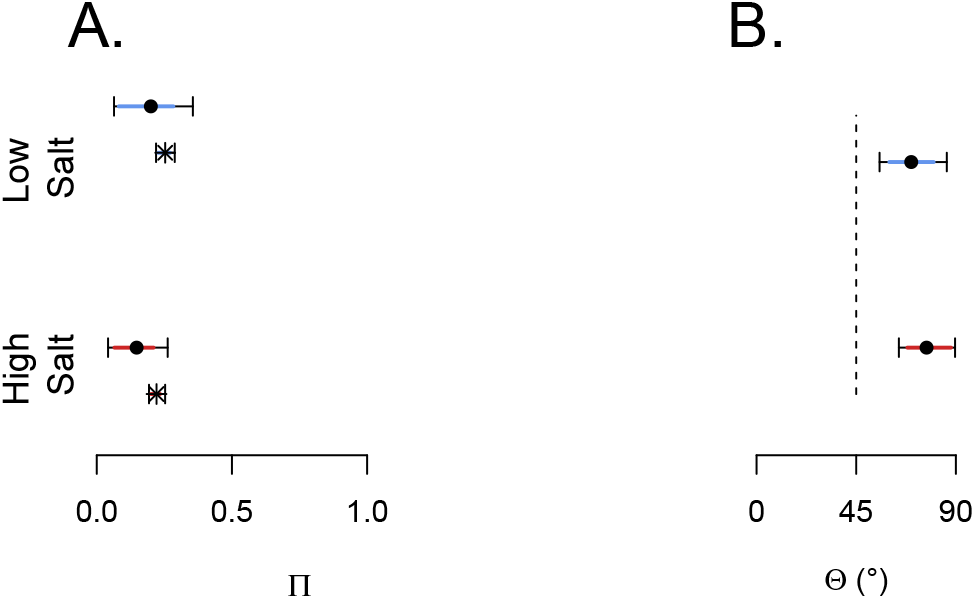
Plasticity and *g*_*max*_ in the ancestral population. **A**. Projection of the amount of genetic variance along the phenotypic plasticity vector Δ*P*. Bars indicate for both low and high salt environments the estimated projections (Π), with dots and error bars the mean, 80% and 95% credible intervals (see Methods). Lower bars show the null expectation if the total genetic variance was equally distributed along all the phenotypic dimensions (round **G**-matrix). **B**. The angle between Δ*P* and *g*_*max*_, Θ, in both low and high salt is higher than the random expectation of 45°C (dashed line, see Methods). Bars indicate 80% and 95% credible intervals.

Phenotypic plasticity (Δ*P*) is not aligned with *g*_*max*_ in the high salt environment, because the amount of genetic variance along phenotypic plasticity is not different than expected by chance (Figure 3A, see Methods), and because the angle between the two phenotypic dimensions of plasticity and *g*_*max*_ is larger than expected (Figure 3B).

### 4.2 Evolution of locomotion behavior in changing salt environments

Three replicate populations from A6140 were exposed to a gradually increasing salt concentration for 35 generations followed by 15 generations in the target high salt environment (GA[1,2,4] populations, see Methods). In the high salt environment, after 50 generations, we find that for each of the transition rates there was at least one replicate population different from the ancestor (Table 3). Among replicates, significant phenotypic divergence for four out of the six transition rates is also observed (Figure 4, Table 4). Phenotypic divergence was not limited by the amount of genetic variance, as the backward to still and forward to still transition rates diverged while showing the lowest genetic variance in the ancestral population (Figure 2). In the low salt environment, there was phenotypic divergence for 3 out of the 6 transition rates (Supplementary Figure S2, Tables 3 and 4).

**Table 3:** Divergence of each GA population from the ancestral population

| transition rate | Low Salt |  |  |  |  |  | High Salt |  |  |  |  |  |
| --- | --- | --- | --- | --- | --- | --- | --- | --- | --- | --- | --- | --- |
|  | divergence |  |  | P value | Chisq | df | divergence |  |  | P value | Chisq | df |
|  | GA150 | GA250 | GA450 |  |  |  | GA150 | GA250 | GA450 |  |  |  |
| SF | -0.0036 | 0.21 | 0.012 | 6.3 e-04 | 17.2 | 3 | -0.21 | 0.045 | -0.048 | 5.9 e-04 | 17.4 | 3 |
| SB | 0.095 | 0.27 | 0.25 | 7.5 e-05 | 21.7 | 3 | -0.16 | 0.18 | 0.13 | 6.3 e-06 | 26.9 | 3 |
| FS | -0.085 | -0.086 | -0.098 | 0.014 | 10.67 | 3 | -0.032 | -0.033 | -0.061 | 0.48 | 2.49 | 3 |
| FB | -0.19 | -0.34 | -0.035 | 9.6 e-05 | 21.1 | 3 | -0.35 | -0.51 | -0.15 | 4.4 e-09 | 41.8 | 3 |
| BS | -0.085 | -0.11 | -0.14 | 1.9 e-06 | 29.3 | 3 | -0.073 | -0.085 | -0.15 | 1.8 e-07 | 34.2 | 3 |
| BF | -0.28 | -0.24 | -0.20 | 7.2e-04 | 16.9 | 3 | -0.42 | -0.52 | -0.31 | 1.9 e-11 | 52.9 | 3 |

**Figure 4:**
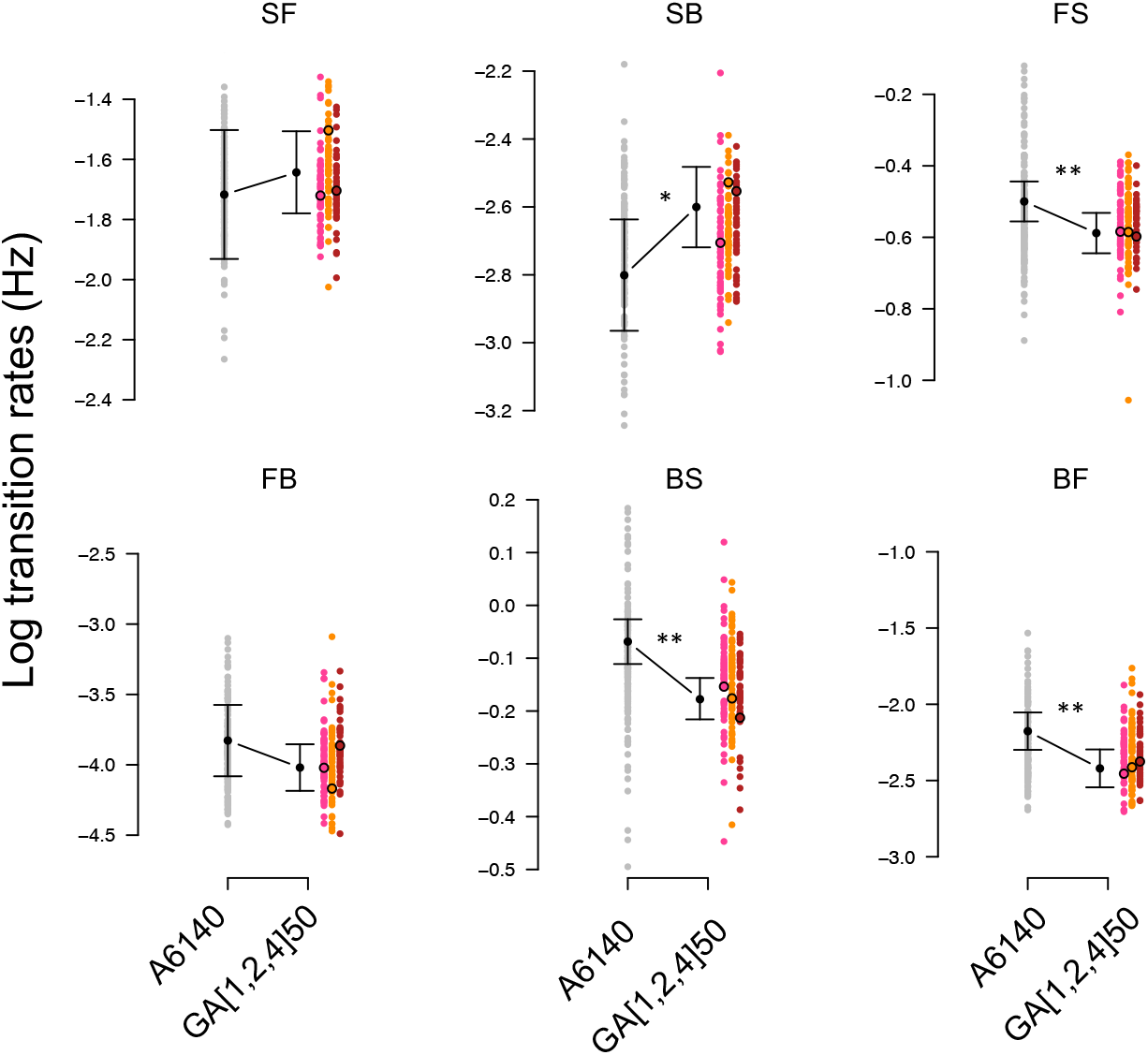
Experimental evolution of locomotion behavior. Small colored dots show the estimated transition rates in high salt for each line from the ancestral population A6140 (grey) and three replicate GA[1,2,4] populations at generation 50 (red). Least square estimates of the mean and 95% confidence intervals are shown in circles and error bars. Divergence between ancestral and the 3 replicate populations is shown by asterisks (LRT *χ*^2^, P-value <0.05 *, <0.01 **, <0.001 ***; Table 4). Divergence per replicate population can be found in Table 3. Results obtained in the low salt environment can be found in Supplementary Figure S2, and analysis in Tables 3 and 4.

**Figure 5:**
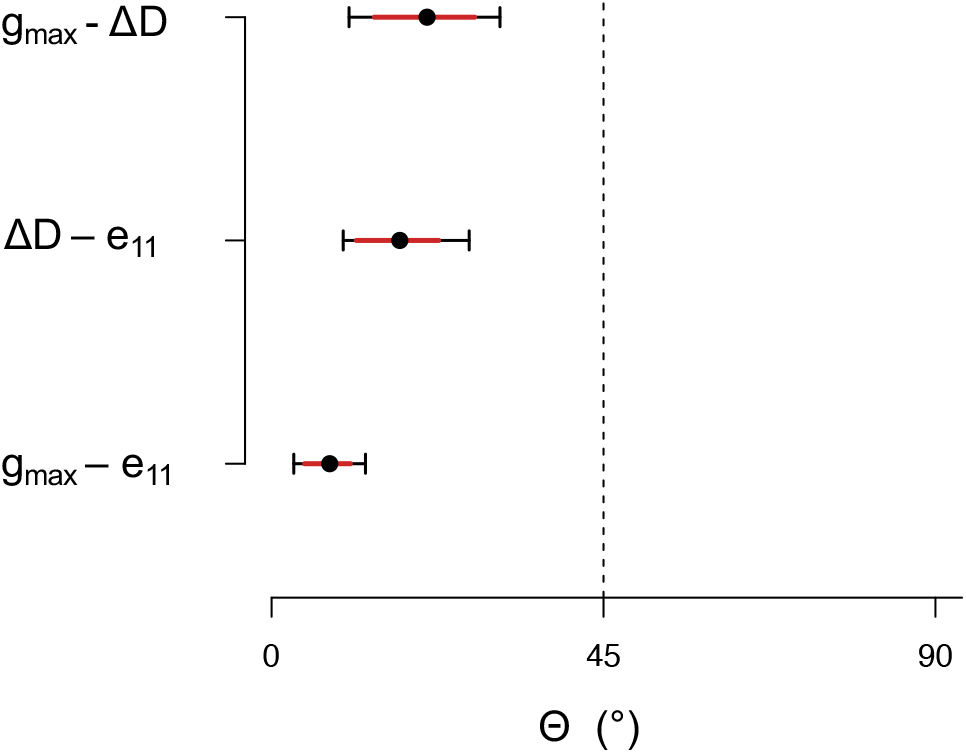
Ancestral *g*_*max*_ with phenotypic and genetic divergence in high salt. The among replicate GA populations angle between ancestral *g*_*max*_ (Figure 2C) and phenotypic divergence Δ*D* (equation 2). Similarly the angle between phenotypic divergence Δ*D* and genetic divergence (*e*_11_, Supplementary Figure S4, and between *g*_*max*_ and *e*_11_. All bars indicate with dots and error the mean, 80% and 95% credible intervals obtained by sampling the posterior distribution estimates of the A6140 **G** matrix (see Methods). The null expectation of non-alignment between vectors is 45° (see Methods), thus the dimension encompassing most genetic variation in the ancestral population aligns with both phenotypic and genetic divergence observed after evolution to high salt.

**Figure 6:**
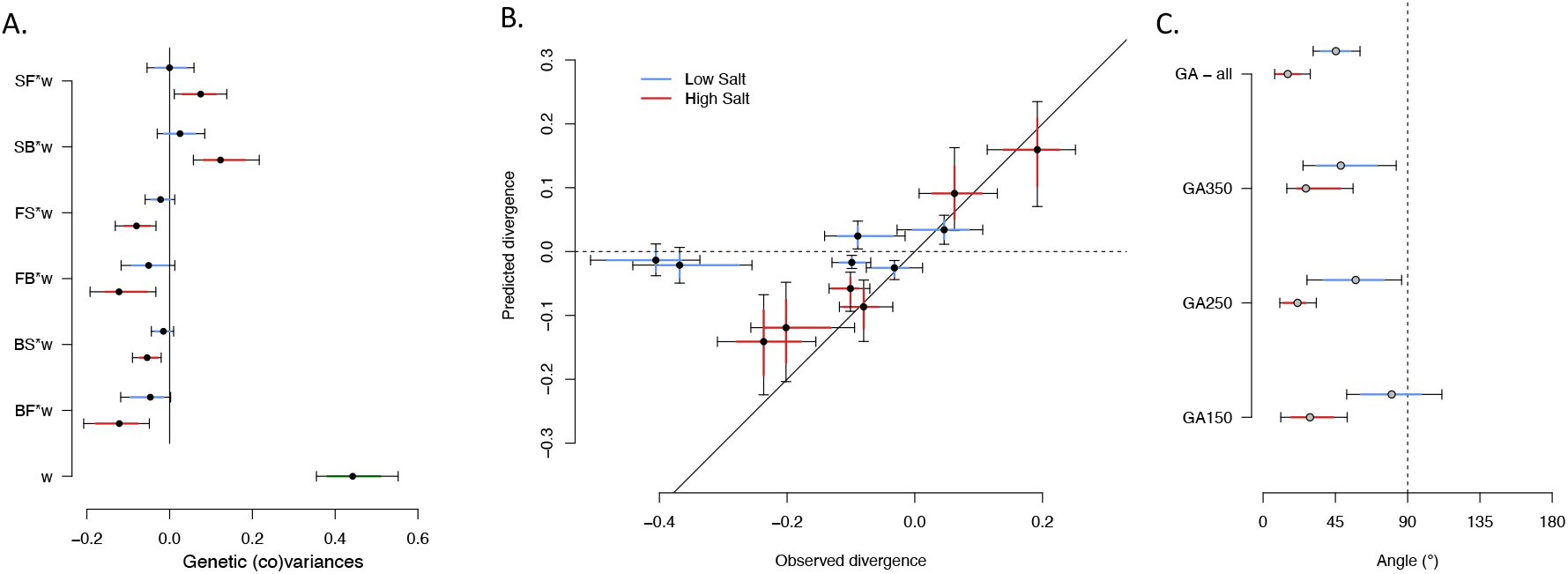
Predicted versus observed phenotypic divergence. **A**. Shown are the ancestral genetic covariances between transition rates in low (blue) and high (red) salt environments with relative fertility in high salt (Robertson’s selection differentials *s* or selection responses *R*, see equation 7). Genetic variance for fitness in high salt is also important (bottom bar). **B**. Shows selection differentials *s* (from panel **A**.) against the observed phenotypic divergence Δ*D* (equation 2, from Figure 4 and Supplementary Figure S2). While in high salt the direction of divergence for all six transition rates are well predicted, because they fall within the identity line, in low salt the direction of divergence are well predicted for only two transition rates (dashed lines showing no predicted divergence). **C**. from panel **B**. we plot the angles between expected and observed phenotypic divergence vectors. For all panels, dots and error bars indicate the mean and the 80% and 95% credible intervals (see Methods). See Supplementary Figure S8 for the predicted versus observed magnitude of phenotypic divergence.

**Table 4:** Divergence among GA populations from the ancestral population

| transition rate | Low Salt |  |  |  |  | High Salt |  |  |  |
| --- | --- | --- | --- | --- | --- | --- | --- | --- | --- |
|  | GA[1,2,4]50 divergence | P value | Chisq | df |  | GA[1,2,4]50 divergence | P value | Chisq | df |
| SF | -0.073 | 0.51 | 0.47 | 1 |  | 0.074 | 0.44 | 0.59 | 1 |
| SB | 0.049 | 0.74 | 0.11 | 1 |  | 0.20 | 0.038 | 4.3 | 1 |
| FS | -0.040 | 0.15 | 2.1 | 1 |  | -0.088 | 0.0083 | 7.0 | 1 |
| FB | -0.34 | 0.050 | 3.8 | 1 |  | -0.19 | 0.13 | 2.2 | 1 |
| BS | -0.10 | 0.019 | 5.5 | 1 |  | -0.11 | 0.0034 | 8.6 | 1 |
| BF | -0.42 | 0.0027 | 9.0 | 1 |  | -0.24 | 0.0045 | 8.1 | 1 |

We next estimated the **G** matrices of derived populations to find about the main phenotypic dimensions of genetic divergence after 50 generations. After 50 generations of evolution there was an overall reduction in genetic (co)variances (Supplementary Figure S3). The three replicate GA[1,2,4] populations mostly diverged from the ancestral population along only one dimension, as defined by the first eigenvector of the first eigentensor measuring the genetic differences among all populations (called here *e*_11_, see Methods; Supplementary Figures S4 and S5).

The phenotypic dimension which contains most genetic variation in the ancestral A6140 population (*g*_*max*_) aligns with the dimension of phenotypic divergence (Δ*D*) in the the target high salt environment, as the angle between both vectors is small (Figure 5). There was consistent parallel genetic divergence of the three replicate populations because the main dimension of genetic divergence (*e*_11_) between them and the ancestor also aligns with the *g*_*max*_ of the ancestor. As a consequence, phenotypic divergence (Δ*D*) and genetic divergence (*e*_11_) follow similar directions. As expected from the analysis in Figure 3, phenotypic divergence did not follow ancestral phenotypic plasticity (Supplementary Figure S6).

### 4.3 Selection and locomotion behavior divergence

Robertson’s selection differentials could predict phenotypic divergence in the target high salt because the are defined as the genetic covariances between traits and relative fitness (see Introduction). These genetic covariances were jointly estimated with the matrix **G**_*qw*_, combining the inbred line transition rates and relative fertility means for the ancestral population (see Methods). We assume that fertility is a good proxy for fitness.

The **G** matrix presented above is robust to the **G**_*qw*_ matrix estimation (Supplementary Figure S7). Transition rate covariances with fertility, for example, reflect the modularity found in the **G**-matrix (Figure 2), as they show signed covariances with fertility similar as those between each other. In high salt, the six transition rate covariances with fertility are different from zero, while in low salt they are not (Figure 6A). Fitness variance in the ancestral population at the high salt environment is high, unlike in the low salt environment (Mallard et al., 2019), confirming that the domestication to the low salt environment and that the high salt environment is a “stressful” environment (Figure 6A).

For all transition rates in high salt, Robertson’s expected responses to selection (*R*) match well with the observed phenotypic divergence (Δ*D*) as shown in Figure 6B. We further computed the angles between the predicted and observed phenotypic divergence vectors and also find a good match between them (Figure 6C). But while in high salt the direction of phenotypic divergence is predicted for all 6 transition rates, its magnitude was not (Supplementary Figure S8): across all populations, observed divergence is only on average 1.3 times the expected per-generation divergence. In the low salt environment predictions for the direction of phenotypic divergence degenerate (Figure 6BC).

Direct selection on transition rates, as measured by the genetic directional selection gradients (*β*_*g*_), were computed as the inverse of the ancestral **G**-matrix obtained from the **G**_*qw*_ matrix and the vector of selection differentials (*s*). We find that *β*_*g*_ are not different from zero, except perhaps for the transition rate between still and backward where it may be positive (Figure 7). From the **G**_*qw*_ modeling we also obtained the vector of selection differentials between transition rates in low salt and fertility in high salt. Unsurprisingly, *β*_*g*_ on low salt transition rates are not different from zero (not shown).

**Figure 7:**
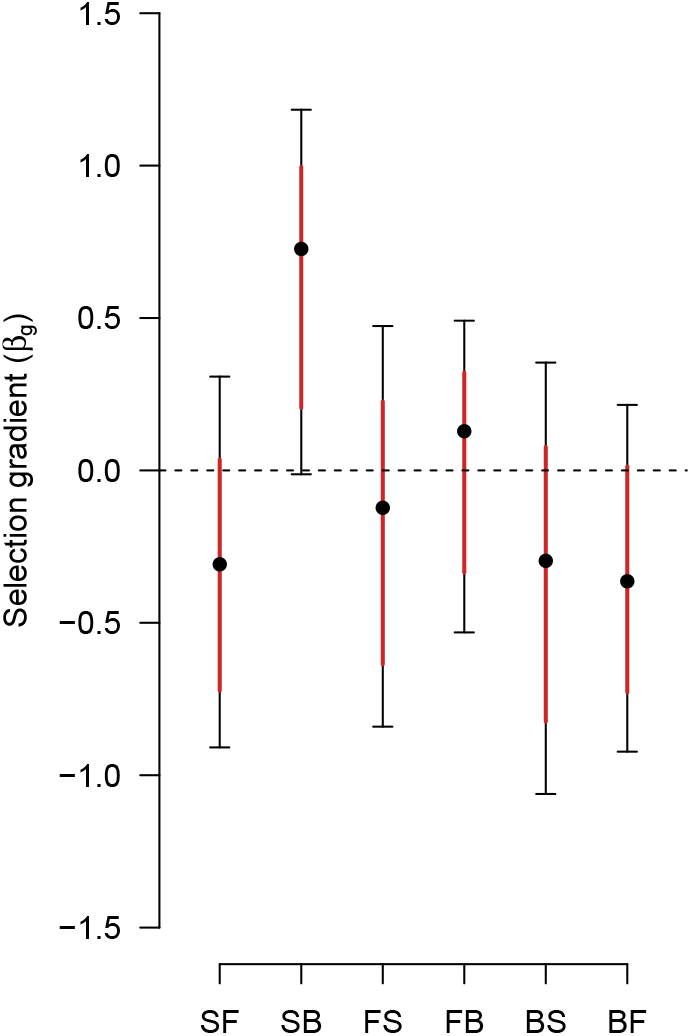
Selection gradients on transition rates. Directional selection gradients on transition rates in high salt are show following equation 8, Lande’s retrospective equation, as the product of the inverse of ancestral **G** matrix in high salt (Figure 2) with Robertson’s selection differentials (equation 7, Figure 6A). Dots and error bars show the mean and 80% and 95% credible intervals obtained by sampling the posterior distribution of the ancestral **G** matrix (see Methods). Similar results are found when the ancestral **G** matrix in low salt are sampled (not shown).

## 5 Discussion

We did experimental evolution with partially-outcrossing *C. elegans* populations harboring standing genetic variation in a gradually increasing salt environment for 35 generations followed by 15 generations in constant high salt. We have previously shown that these partial-outcrossing populations exposed to changing salt environments purged deleterious recessives alleles while maintaining genetic diversity during the 50 generations of evolution, either due to associative overdominance or true overdominance (Chelo et al., 2019). The derived populations adapted to the high salt conditions (Theologidis et al., 2014), and thus here we could ask: 1) if plasticity in the transition rates between movement and direction of locomotion behavior were aligned with the phenotypic dimension where more genetic variation is found in the ancestral population and, as a consequence, help explain adaptive phenotypic divergence in high salt; 2) whether or not 50 generations of phenotypic divergence could be predicted from the ancestral genetic covariances of transition rates with fitness in high salt, using Robertson’s secondary theorem of natural selection; and 3) if direct and/or indirect selection on transitions was responsible for phenotypic divergence, using Lande’s equation with Robertson’s selection differentials. For an analysis of selection on locomotion behavior, and the evolution of the **G** matrix, in a constant low salt environment we refer the reader to Mallard et al. (2019).

Fifty generations of experimental evolution in changing salt conditions led to phenotypic divergence in the target high salt environment for at least 5 out of the 6 transition rates, and in the low salt environment for 3 out of the 6 transitions rates, although there was heterogeneity between replicate populations. In the ancestral population, and in both salt environments, transition rates between still and forward (SF) and still and backwards (SB) form different behavioral modules as these traits show positive genetic covariances with each other, and negative genetic covariances with all other transition rates. Gene expression in our populations is restricted to being active (leaving the still state) and not the direction of movement. In itself this observation is interesting because it explains why phenotypic divergence in SF and SB is of different sign than that of the other 4 transition rates, and because it also explains why the phenotypic dimension encompassing most genetic variation of the ancestral population aligns well with the main dimensions of both phenotypic divergence and genetic divergence among all populations in high salt. However, many studies of neuronal, muscular or metabolic function using mutants have suggested that engaging in activity can be decoupled from the direction of movement (De Bono and Maricq, 2005; Zhen and Samuel, 2015) and elsewhere we have described little mutational pleiotropy for transition rates for two strains founders of our populations (Mallard et al., 2022). The evolution of genetic covariances between transition rates in our populations should therefore result mostly from the frequency dynamics of relevant alleles in linkage disequilibrium (Lande, 1980; Phillips and McGuigan, 2006). See Mallard et al. (2022) for a discussion on the role of pleiotropy and linkage disequilibrium in some of our populations.

Several authors have suggested that ancestral phenotypic plasticity will be aligned with the phenotypic dimensions with more standing genetic variation, or mutational input, and thus facilitate adaptation if past correlated selection between traits led to the evolution of plasticity in particular dimensions of trait combinations (Arnold et al., 2008; Draghi and Whitlock, 2012; Jones et al., 2007, 2014). Empirical tests of this idea are few. In a meta-analysis of several studies in natural populations phenotypic plasticity generally matched phenotypic dimensions with more genetic variation but did not depend on the degree of environmental stress and thus, presumably, on selection (Noble et al., 2019). More explicitly, Johansson et al. (2021) have showed that phenotypic plasticity for life-history and morphological trait responses to changes in photoperiod in damselflies was aligned with phenotypic dimensions with more genetic variation but only in populations facing selection because of photoperiod seasonality, as expected from theory. In our study, the **G** matrix is very similar in the low and high salt environments for the ancestral population, and the phenotypic dimension with more genetic variation in high salt neither aligns well with that of phenotypic plasticity nor the phenotypic dimension where plasticity is found contains much genetic variation. It is thus not surprising that the mismatch between plasticity and genetics does not impede or facilitate adaptive phenotypic divergence.

One of the main results of our study is that the direction of phenotypic divergence for at least 5 out of 6 transition rates in high salt is predicted using Robertson’s equation. The direction of divergence in the transition rate between still and forward is not predicted, even if it has a detectable genetic covariance with fitness because of replicate population heterogeneity and perhaps a founder sampling effect when establishing them. Yet, these are remarkable findings, especially when considering that the individual selection surface likely changed during the first 35 generations of the experiment, and that the **G** matrix in high salt evolved during the 50 generations, by drift and selection, cf. (Guzella et al., 2018), mostly through a reduction in genetic variance across all phenotypic dimensions. The selection surface probably changed during evolution but only in strength, because only one environmental factor was employed and the nature of change across generations was linear on the mean environmental value, not the variance (Chevin et al., 2015; Proulx and Teotónio, 2017). That the Robertson equation predicts phenotypic divergence is also remarkable because fertility is only a component of fitness. Indeed, we previously were unable to measure increased fertility in high salt (Chelo et al., 2019), despite obvious adaptation – as measured with competition assays between ancestral and derived populations (Teotónio et al., 2017), see Figure 4d in Theologidis et al. (2014) –. Our analysis demonstrates that even if we estimated broad-sense genetic covariances between transition rates and fertility they are nonetheless adequate surrogates of the additive genetic covariances of outbreeding populations, as required by Robertson’s equation.

But while characterization of genetic covariances between transition rates and fertility in high salt in the ancestral population is sufficient to predict the direction of phenotypic evolution in the target high salt environment, we cannot predict with much accuracy the direction of phenotypic evolution in the (ancestral) low salt environment, and hence we cannot predict the evolution of phenotypic plasticity. The simplest explanation for this finding is that the ancestral population contained low and significant levels of genetic covariances between transition rates in low salt and fitness in high salt, but that these covariances were undetectable given sample sizes we have. In low salt, transition rates are effectively neutral and could have only diverged by being genetically correlated with high salt traits under selection. Similarly, in Mallard et al. (2019) we have described that phenotypic stasis of locomotion behaviour in populations facing constant low salt results from effective stabilizing selection over a large phenotypic space after adaptation, cf. (Haller and Hendry, 2014), and that after adaptation, Robertson’s equation of little use to predict evolution over local phenotypic space because in this local space genetic covariances between transition rates and fitness are small and mostly evolve by drift (Mallard et al., 2019).

The Robertson equation has been applied to a few natural populations, those where time series of trait and fitness values are concurrently obtained with pedigrees such that trait breeding values can be estimated. Biquet et al. (2022) provide an exemplary study of a blue tit population that has been followed for more than 40 years, and in which egg laying has changed to earlier and earlier spring dates, presumably because of climate change. Despite significant heritability for egg laying date and selection on it, however, modeling the expected breeding values for laying date did not reveal any temporal trend consistent with a general lack of genetic covariance between laying date and fitness during the whole period followed. There was thus no evolution, meaning no genetic divergence for laying date, which the authors can attribute to the stochastic nature of individual development to maturity. Using a similar approach, Bonnet et al. (2017) have found on the contrary that evolution towards earlier dates of parturition in a red deer population (also over 40 years and also presumably because of climate change) could be predicted and were consistent with the estimated change of parturition date breeding values. What our study suggests is that for organisms where pedigrees are difficult to obtain, and thus trait breeding values difficult to estimate, such as small organisms that cannot be marked and those with a rapid or complex life-cycle, for example, one may be able to measure the genetic covariance of the traits of interest with fitness in an ancestral population, to predict the direction of adaptive future phenotypic divergence on the short-term of a few tens of generations.

A second major finding of our study is that we were unable to infer direct selection on transition rates, despite phenotypic divergence in a majority of them, as directional selection gradient estimates were not obviously different from zero. There are two explanations for this result: either unobserved traits under direct selection are genetically correlated with observed ones or there is correlated selection between observed and unobserved traits because of unknown environmental covariances of them with fitness. A recent study in wild salmon as shown that selection due to human harvesting of a prey species (for feeding cultured salmon) explained divergence for early maturity and small body sizes, despite selection for increased body size at maturity because of fishing (Czorlich et al., 2022). In an example privileging the second explanation, Kruuk et al. (2002) have found that despite heritability and directional selection for antler size in red deer there was phenotypic stasis due to nutritional state covariation and antler size. We find it unlikely, however, that a similar situation to Kruuk et al. (2002) explains our results. This is because worms’ locomotion behaviour were measured in ancestral and derived populations in “common garden” assays, after reviving population samples from storage and culturing them under similar environmental conditions for 4-6 generations. Perhaps more importantly, we have also reported that transgenerational environmental effects on the viability of individuals exposed to high salt do not carry over for more than three generations, and are in any case small when compared to the fitness effects of within-generation high salt exposure (Proulx et al., 2019). Expression of trait values specific to changing salt environments could be at play, but this would require a genetic basis for individuals to detect and anticipate the higher salt levels of future generations, despite they themselves developing and growing under lower and constant salt levels as was the case during experimental evolution (Dey et al., 2016; Proulx and Teotónio, 2017).

We favor the interpretation that indirect selection on transition rates is due to them being genetically correlated with unobserved traits under direct selection. One candidate trait to be under direct selection is body size at the time of reproduction. In a changing salt environment increased body size could be favored for example because the accumulation of osmolites will lower cuticle tension and enhance individual survival. Body size and locomotion behavior in *C. elegans* are further known to be physiologically regulated in concert not only by the amount of food eaten, and how much energy is spent by growing, foraging, mating or reproducing, but also by the perception of stressful environments (Fujiwara et al., 2015, 2002). The software tracker that we used to obtain locomotion behavior data also contains information about body size (Noble et al., 2017; Swierczek et al., 2011). Estimates of the genetic covariance between body size and fertility in the ancestral population suggest an expected response to selection in high salt of 0.2, which is similar to the observed body size change after 50 generations of evolution and, coincidentally, a selection gradient of *β*_*g*_ = 0.2 (−0.03-0.46, 95% CI). We also find significant genetic covariances between all transition rates and body size (not shown). Many other traits can be implicated, however, for example developmental time, because hermaphrodites engaged in selfing tend to more rapidly resume activity when harassed by males than when self-sperm depleted (Chasnov et al., 2007; Lipton et al., 2004) and delayed maturity favors selfing in this protandrous nematode (Poullet et al., 2016; Theologidis et al., 2014). Only a more detailed investigation can shed light on the complexity of the selection surface in the high salt environment. Our analysis suggests, however, that indirect selection led to the divergence of locomotion behavior in part because of important genetic covariances between body size and transition rates.

In our experimental evolution model, ancestral plasticity neither hinders nor facilitates phenotypic divergence of locomotion behavior. The direction of phenotypic divergence in the target environment can be predicted from the ancestral genetic covariances of component traits of locomotion behavior with fitness, even if experimental evolution took place in a gradually changing environment until reaching the target environment, for multiple generations, and there was a large reduction of genetic variation affecting locomotion behaviour. Indirect selection seems to be responsible for the phenotypic divergence of locomotion behavior. Together, these findings indicate that selection theory can be used to predict the direction of short-term adaptive phenotypic evolution.

## 6 Acknowledgments

We thank I. Chelo, H. Gendrot, T. Guzella and L. Noble for help with worm handling, data acquisition or data analysis; L. Noble, P. Phillips and P. de Villemereuil for discussion. This work was supported by the European Research Council (ERC-St-243285) and the Agence Nationale pour la Recherche (ANR-14-ACHN-0032-01, ANR-17-CE02-0017-01).

## 7 Author contributions

Data acquisition BA; Software implementation and data analysis FM; funding acquisition and project administration HT; Conceptualization, writing, editing and correspondence: FM, HT.

## 9 Supplementary Figures

**Figure S1:**
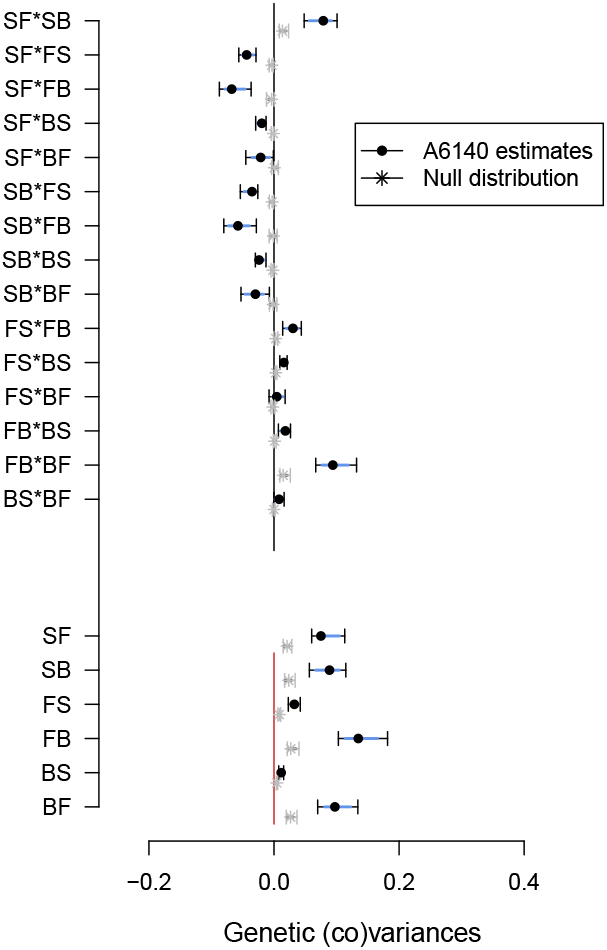
**G**-matrix of the ancestral population A6140. **A. G** matrix estimates in the low salt environment, bottom six entries indicating the genetic variances of transition rates, top 15 entries indicating the the genetic covariances between transition rates. Colored intervals and bars show 80% and 95% credible intervals of the posterior distribution, dots the mean (see Methods). In grey, the expected genetic (co)variances due to sampling a finite number of inbred lines, following Morrissey and Bonnet (2019).

**Figure S2:**
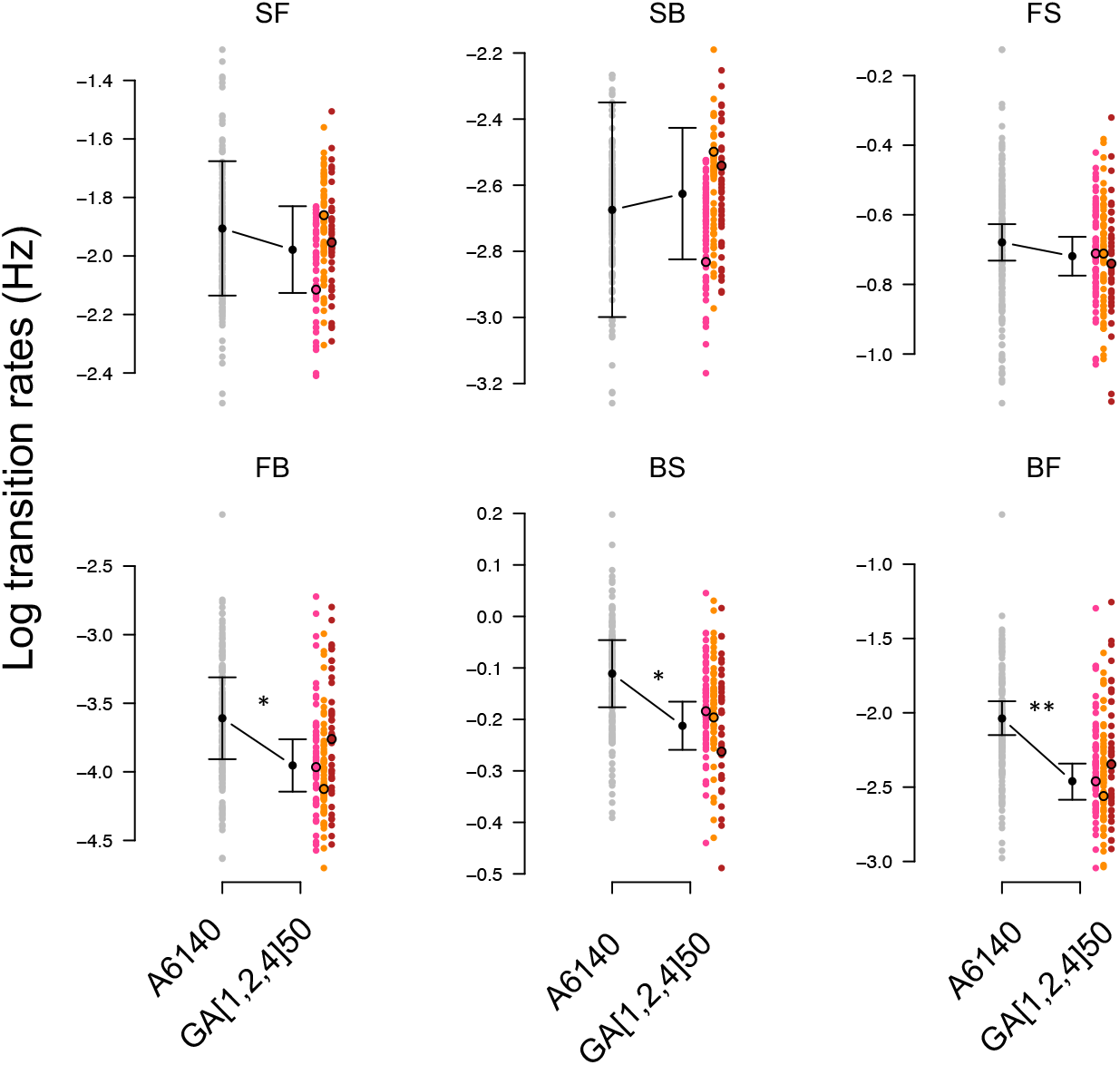
Experimental evolution of locomotion behavior. Small colored dots show the estimated transition rates in low salt for each line from the ancestral population A6140 (grey) and three replicate GA[1,2,4] populations at generation 50 (red). Least square estimates of the mean and 95% confidence intervals are shown in circles and error bars. Divergence between ancestral and the 3 replicate populations is shown by asterisks (LRT *χ*^2^, P-value <0.05 *, <0.01 **, < 0.001 ***; Table 4). Divergence per replicate population can be found in Table 3.

**Figure S3:**
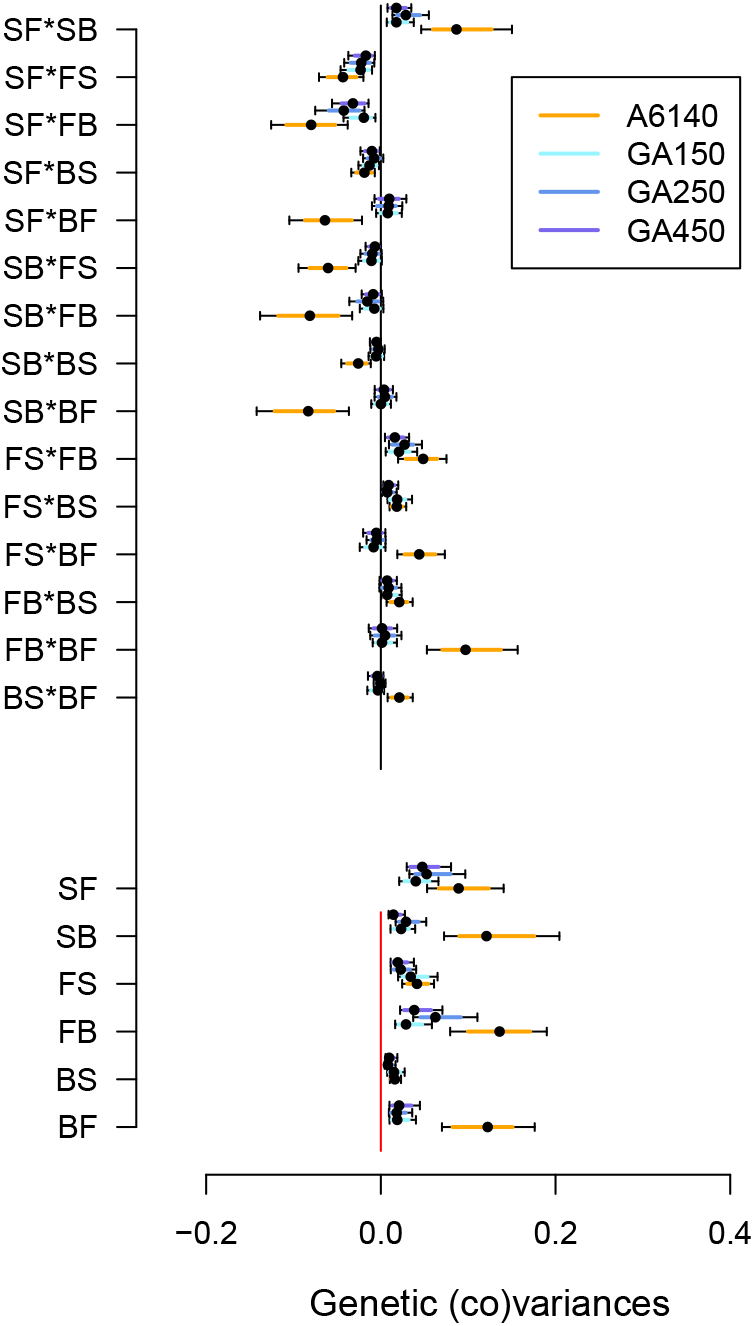
Evolution of the **G** matrix in the high salt environment. Bottom six entries indicate the diagonal estimates the genetic variances of transition rates, top 15 entries indicate the off-diagonal estimates of genetic covariances between transition rates. Orange (blues) indicate estimates of ancestral (derived) populations, with colored bars showing 80% and 95% credible intervals of the posterior distribution, dots the mean.

**Figure S4:**
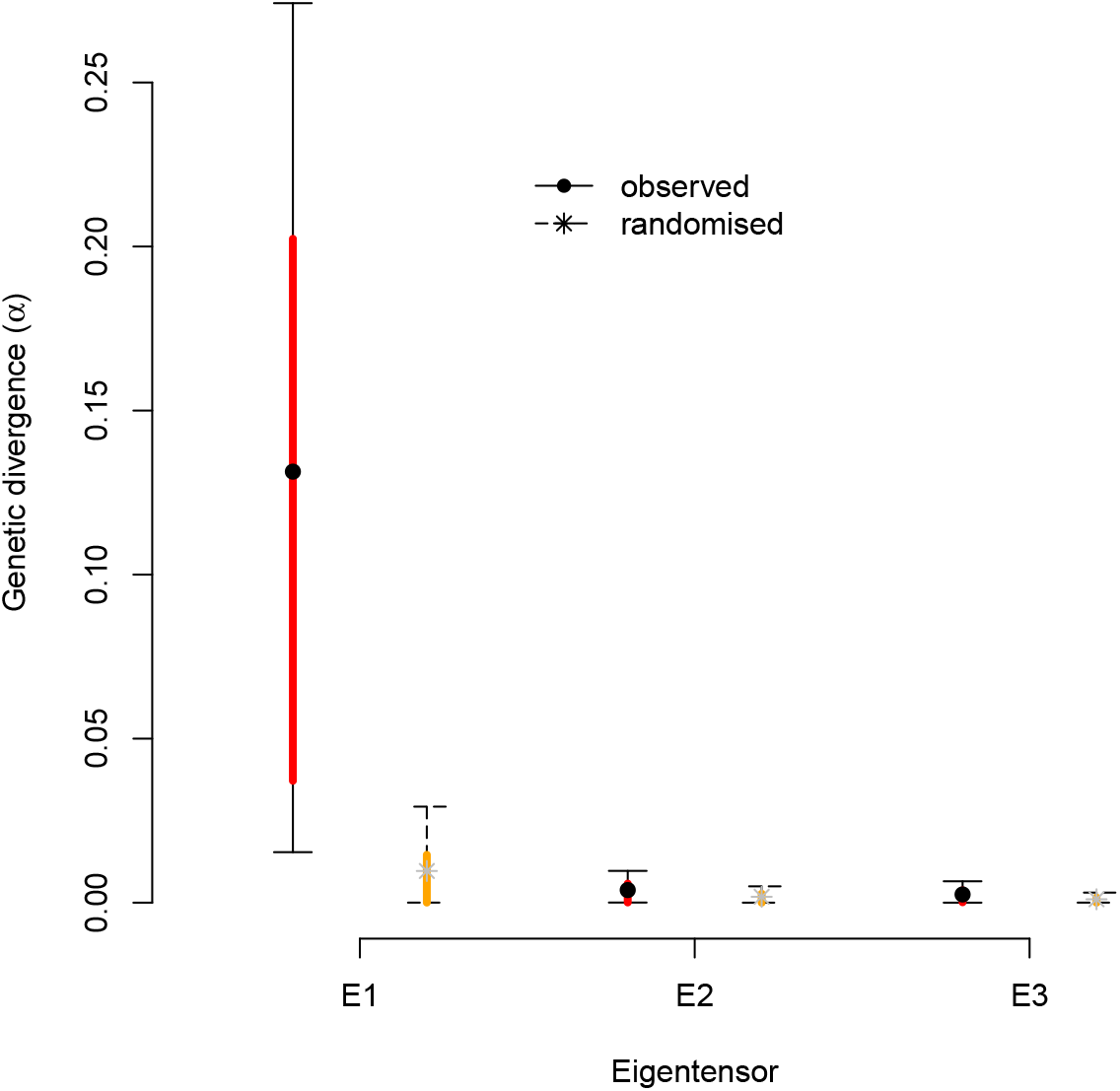
**G** matrix evolution in high salt. **A**. Eigentensor analysis of all four ancestral and derived **G** matrices following Aguirre et al. (2014) and Morrissey and Bonnet (2019). Shown are the first three eigentensor values encompassing most of the differences between **G** matrices, with only the first one *E*_1_ containing more genetic variation than expected with our sample sizes. Colored bars show 80% and 95% credible intervals of the posterior distribution, dots and stars the mean.

**Figure S5:**
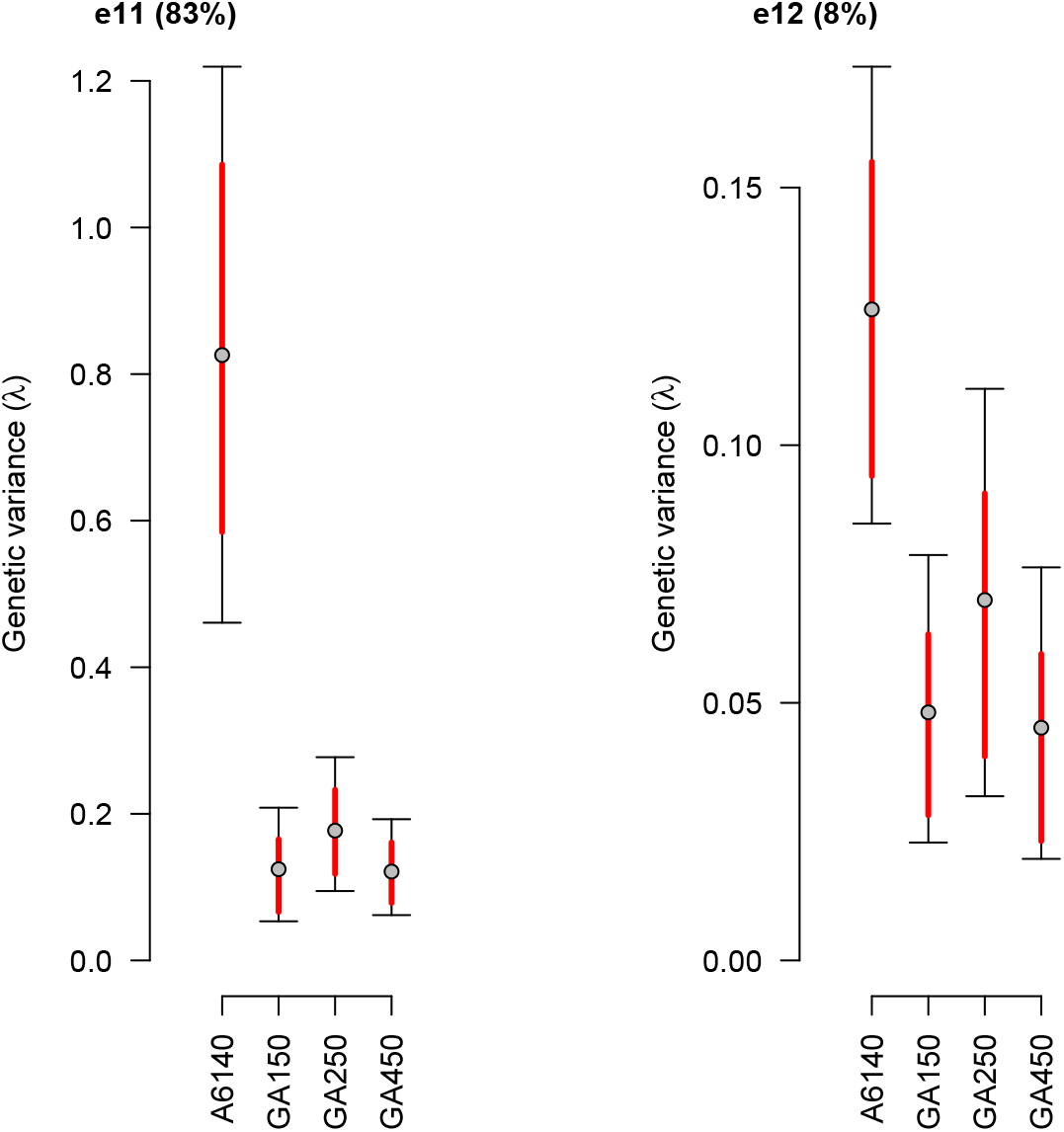
Genetic divergence during evolution. For the first eigentensor *E*_1_ from Supplementary Figure S4B, further decomposition shows that its first eigenvector (*e*_11_) explains 83% of the genetic differences between all populations populations (eigenvalues in y-axis). Colored bars show 80% and 95% credible intervals of the posterior distribution, dots the mean. For the second eigenvector *e*_11_ of the first eigentensor *E*_1_, there was also a reduction of genetic variation between A6140 and GA populations though only accounting for 8% of genetic divergence (not shown).

**Figure S6:**
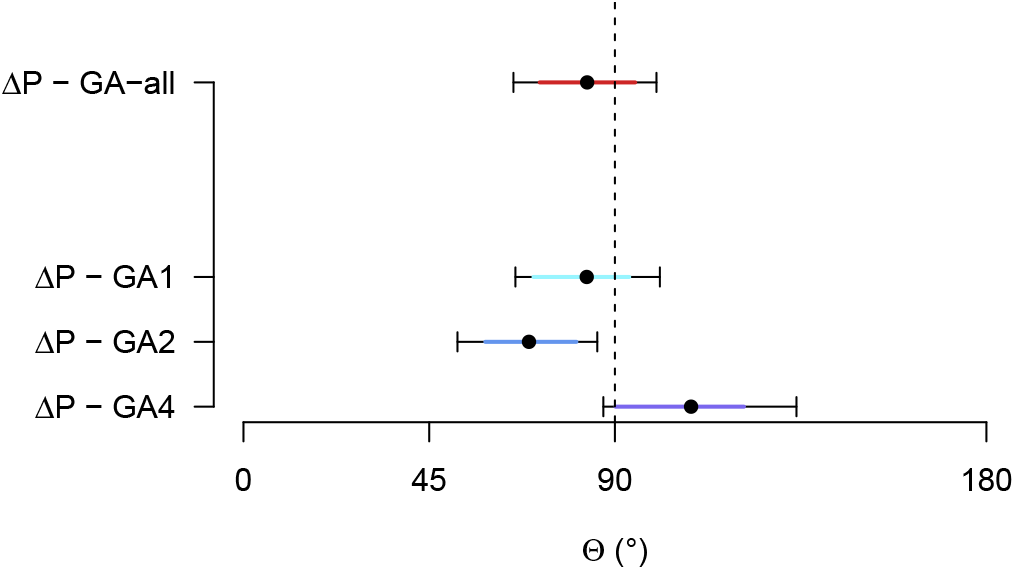
Ancestral phenotypic plasticity and phenotypic divergence. Angles between Δ*P* and Δ*D* among replicate GA populations or for each one of them are shown from top to bottom. Least square estimates of the mean, 80% and 95% confidence interval are represented by dots and colored error bars.

**Figure S7:**
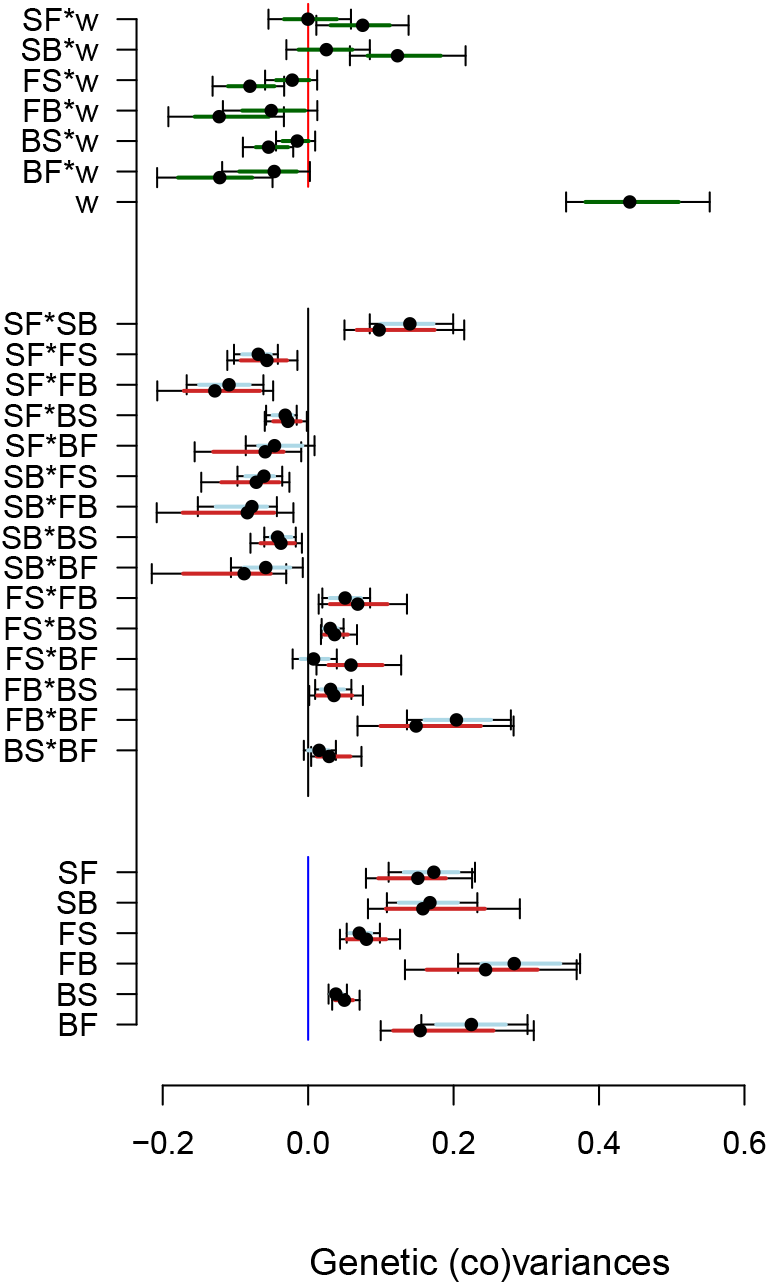
Ancestral *G*_*qw*_ matrix with fitness. *G*_*qw*_ matrix estimates are compared with **G** matrix estimates (Figure 2 and Supplementary Figure S1) for the ancestral population in low (blue) and high (red) salt environments. Bottom six entries indicate the diagonal estimates the genetic variances of transition rates, middle 15 entries indicate the off-diagonal estimates of genetic covariances between transition rates. Top 7 entries indicate the genetic covariances between transition rates and relative fertility or the genetic variance in relative fertility (as in Figure 6). Genetic (co)variances estimates are very similar to those presented in Figure 2B.

**Figure S8:**
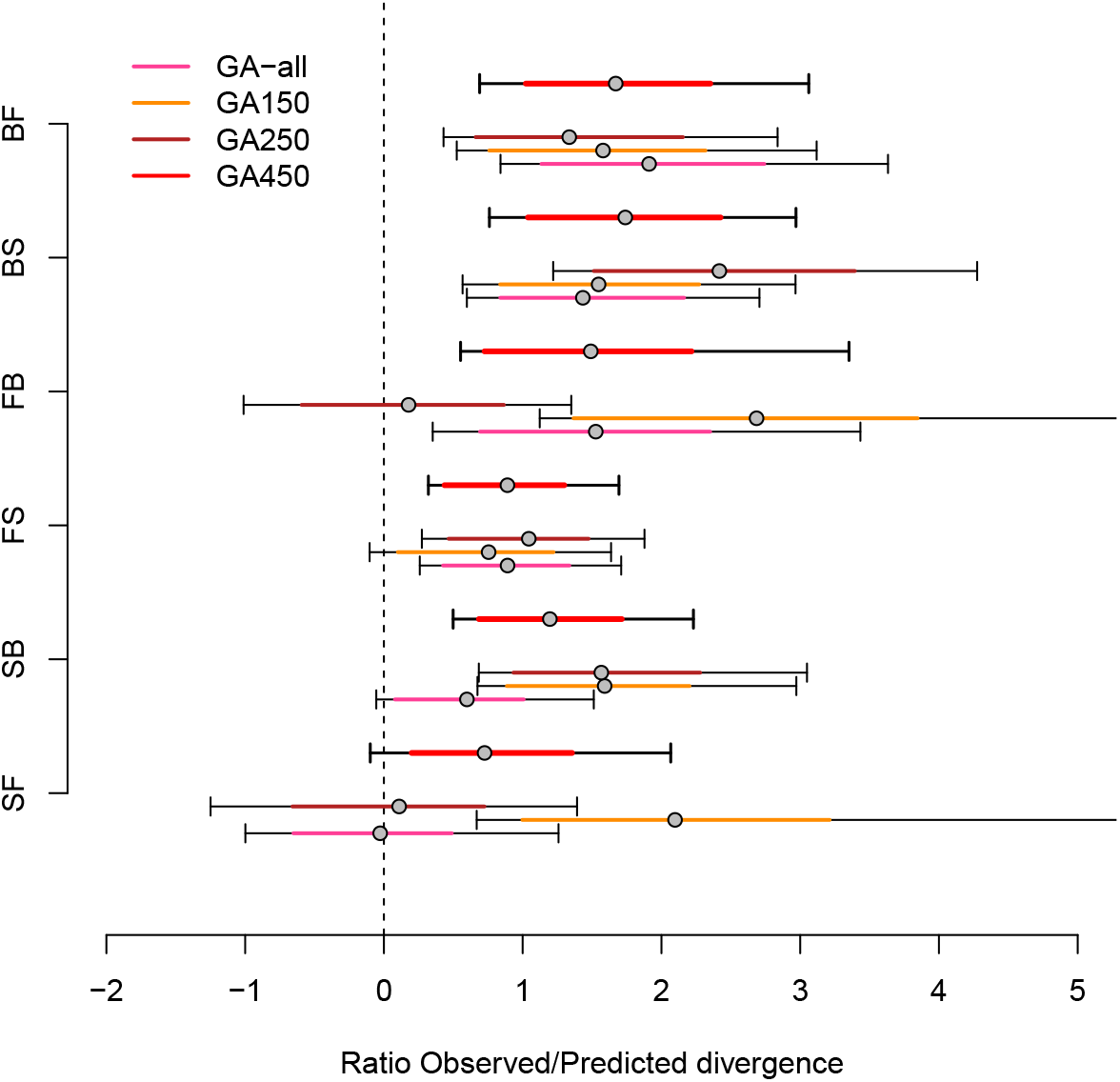
Magnitude of phenotypic divergence. The ratio between the expected phenotypic divergence (*s*, from Figure 6) with the observed phenotypic divergence (Δ*D*) is plotted for all transition rates for each GA replicate. Error bars show 80% and 95% credible intervals, obtained by sampling the posterior distributions; dots the mean. When this ratio falls to zero we cannot predict phenotypic divergence. On average predicted phenotypic divergence would have been achieved in 1.3 generations, much sooner than when GA populations were measured at generation 50.

